# Regularities in species niches reveal the World’s climatic regions

**DOI:** 10.1101/851030

**Authors:** Joaquín Calatayud, Magnus Neuman, Alexis Rojas, Anton Eriksson, Martin Rosvall

## Abstract

Although classifications of the Earth’s climates date back to the ancient Greeks, the climatic regions shaping the distribution of animals remain poorly resolved. Here we present a classification of global climates based on regularities in realised niches of 3657 amphibians, 7204 reptiles, 10684 birds and 4574 mammals. We found 16 main climatic regions that are mostly consistent across groups and previous plant expert-based classifications, confirming the existence of major climatic restrictions for life. The results also suggest that differences among groups likely relate to their particular adaptations and dispersal capabilities. We further show how the integration of species niche classifications with geographical information provides valuable information on potential mechanisms shaping the climatic regions. Our climate classification has applications in several disciplines, including conservation planning and ecological and evolutionary studies.

## INTRODUCTION

Climate governs the basis for life on Earth. Besides historical contingencies and geographical barriers, abiotic conditions determine species ranges [1–3] and derived diversity patterns [4, 5]. On a global scale, distinctive climatic regimes impose generalised restrictions, leading to the formation of species pools adapted to them and ultimately to the generation of biomes [6]. Identifying the boundaries of these climate regimes is, therefore, a fundamental challenge to understand how life organizes on Earth.

Already Pythagoras proposed a classification of climate regimes of the known world in the sixth century BC [7]. However, it was not until the 19th century when geographers laid the foundations for such classifications [8]. By that time, researchers noticed the close relationship between the distribution of various life forms, especially vegetation types, and climate [8]. For instance, Köppen built his long-standing climate classification from pioneer plant classifications, assuming that vegetation forms carry information about climatic conditions [9, 10]. This assumption has received considerable support [11], and the Köppen classification system is widely used nowadays as the standard classification of climates in a range of disciplines, including climatology [12], geography [13], conservation planning [14], and ecology [15]. However, the fact that plant species are good indicators of general climatic conditions does not necessarily imply that such conditions restrict the distribution of other organisms in the same manner. If different taxa have different climatic adaptations, the boundaries defining climate types will vary among them. Following Thornthwaite [10], the “truly active factors” describing a climate type may vary among organisms. Thus, while Köppen’s climate classification can indicate the active climatic factors for plants, it remains unknown whether they are also appropriate for other organisms. Despite several attempts to refine or propose alternative climatic regions [16–19], quantitative studies defining climatic regions for other organisms are still lacking.

The current information on species distributions and global climatic variables, together with recent advances in niche modelling and classification techniques provide an unprecedented opportunity to identify the climatic boundaries shaping the distribution of faunas and floras across the globe. The last decades have witnessed
51 a tremendous collective effort to record occurrences of a large number of species [20], which has resulted in comprehensive datasets with the distributional ranges of several groups [21–23]. Also, data on climatic variables at a global scale have been developed at high spatial resolutions [24, 25]. This information allows to characterise the realised climatic niches of diverse species and to find regularities among them. For example, projecting these realised climate niches into a climatic space [26] should, if climatic boundaries exist, reveal co-occurring groups of species across particular portions of the climatic space. Thus, identifying these portions, or niche domains, should uncover the main climatic boundaries shaping the organization of life (Fig. 1).

**FIG. 1:**
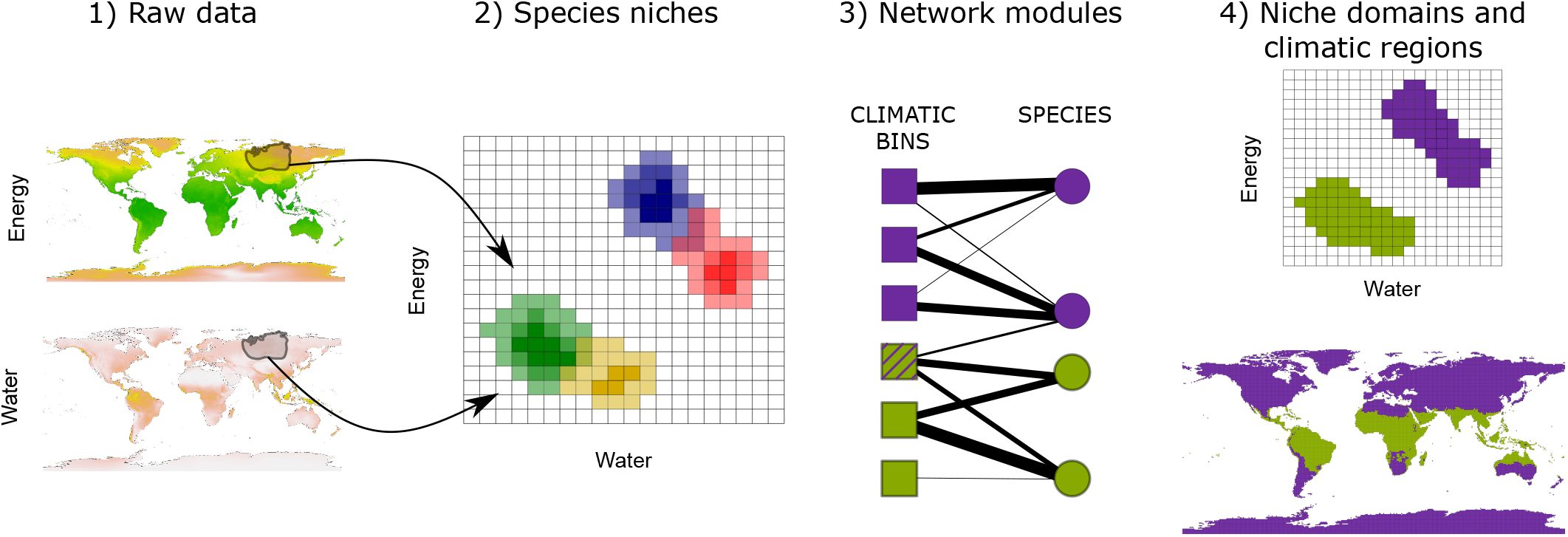
Workow to identify niche domains and climatic regions. Using the climatic conditions a given species experiences within its range (1), we project the species’s niche into a climatic space discretised in an optimal number of bins (Appendix S1) (2). We translate the binned data into a weighted bipartite network, where climatic bins and species form the nodes and the probabilities of finding the species in the bins form the weighted links (3). Using a network community detection algorithm, we identify domains of the climatic space with similar species (4, upper). The climatic conditions defining these domains delineate the corresponding climatic regions of the Earth (4, lower).The striped climatic bin is linked to species classified in both climatic domains and, therefore, it represents a diffuse transition with low specificity.

Besides climate shaping niche domains, dispersal barriers and historical contingencies may also inuence their shape [3, 27, 28]. Therefore, similar climates may have different effects across geographic regions [29]. For instance, while a given climate may lead to specific species pools in some parts of the Earth, the same climate in other parts of the Earth may not hold specific species pools. Such lack of specific species can occur, for example, because the required adaptations have not appeared [30], the adapted species have been not able to disperse [31], or the area is too small to hold large species pools [32]. Thus, studying the signature of these historical and geographical processes, the geographical signal for short, in niche domains can provide valuable information about the potential mechanisms behind them and their associated climatic regions.

Here we explore the global climate regions of Tetrapoda by characterising the climatic niche domains of amphibians, birds, mammals and reptiles. Tetrapoda is a well-suited group for our purpose. First, comprehensive databases are available, including the distributional ranges of most species in the group [21–23]. Second, the different classes within Tetrapoda possess diverse capabilities to disperse and withstand abiotic conditions. Therefore, we can investigate if various capabilities influence climatic niche domains, and possibly generalise the climatic regions to other groups. Third, there is accumulated evidence on the main climatic factors controlling the distribution of these species, which simplifies the selection of appropriate climatic variables. In particular, the distribution of tetrapods is strongly determined by water and energy aspects of climate [4, 33–37]. Finally, researchers study Tetrapoda species in several disparate fields - from animal husbandry [38] to ecological [39] and evolutionary studies [40] – where a description of their climatic regions can be especially useful.

In our classification approach, we first approximate the realised niche of each species as the probability of finding the species across a two-dimensional space that representing water and energy aspects of climate (Fig. 1). We then use a community-detection algorithm from network theory to simultaneously find portions of the climatic niche space holding similar species, the niche domains, and the species grouped into these domains. Mapping back to the Earth’s surface gives for each climatic niche domain a climatic region. We then examine the transition zones and the geographical signal in the climatic regions. The novel climatic regions confirm the existence of generalised climatic constraints across life forms. Therefore, the climatic regions provide valuable information for conservation and ecological and evolutionary studies of Tetrapoda in particular and animals in general.

## RESULTS

### Major climatic niche domains of Tetrapoda

We first identified the niche domains of each Tetrapoda class independently. We calculated the proportion of observations of each species within each bin of a climatic space defined by potential evapotranspiration (PET) and annual precipitation (AP; Fig. 1, Methods and Appendix S1). We represented this data as a weighted bipartite network where climatic bins and species form two disjunct sets of nodes, and the probabilities of finding the species in the bins form the link weights. Using a hierarchical network clustering algorithm [41, 42], we obtained groups of climatic bins holding similar species (i.e. niche domains) and the species most associated with them.

We found similarities among Tetrapoda classes in the detected niche domains, but also observed some differences (Fig. 2). For instance, the number of major domains with 50 or more species in the lower hierarchical level is similar across Tetrapoda classes, ranging from 13 to 15. However, mammals and birds show a domain of low energy, whereas reptiles present some domains across arid conditions, that is with elevated energy inputs and low water availability (Fig. 2). These differences seem to be related to the particular adaptations of each group to withstand climatic conditions. Nevertheless, the classification of most domains was largely congruent across classes, and hence we classified the climatic space of the Tetrapoda superclass by using all species jointly. The niche space of Tetrapoda divided into 16 main domains that were similar to those of the independent classes, and some of the above-explained particularities did not appear (Fig. 2).

**FIG. 2:**
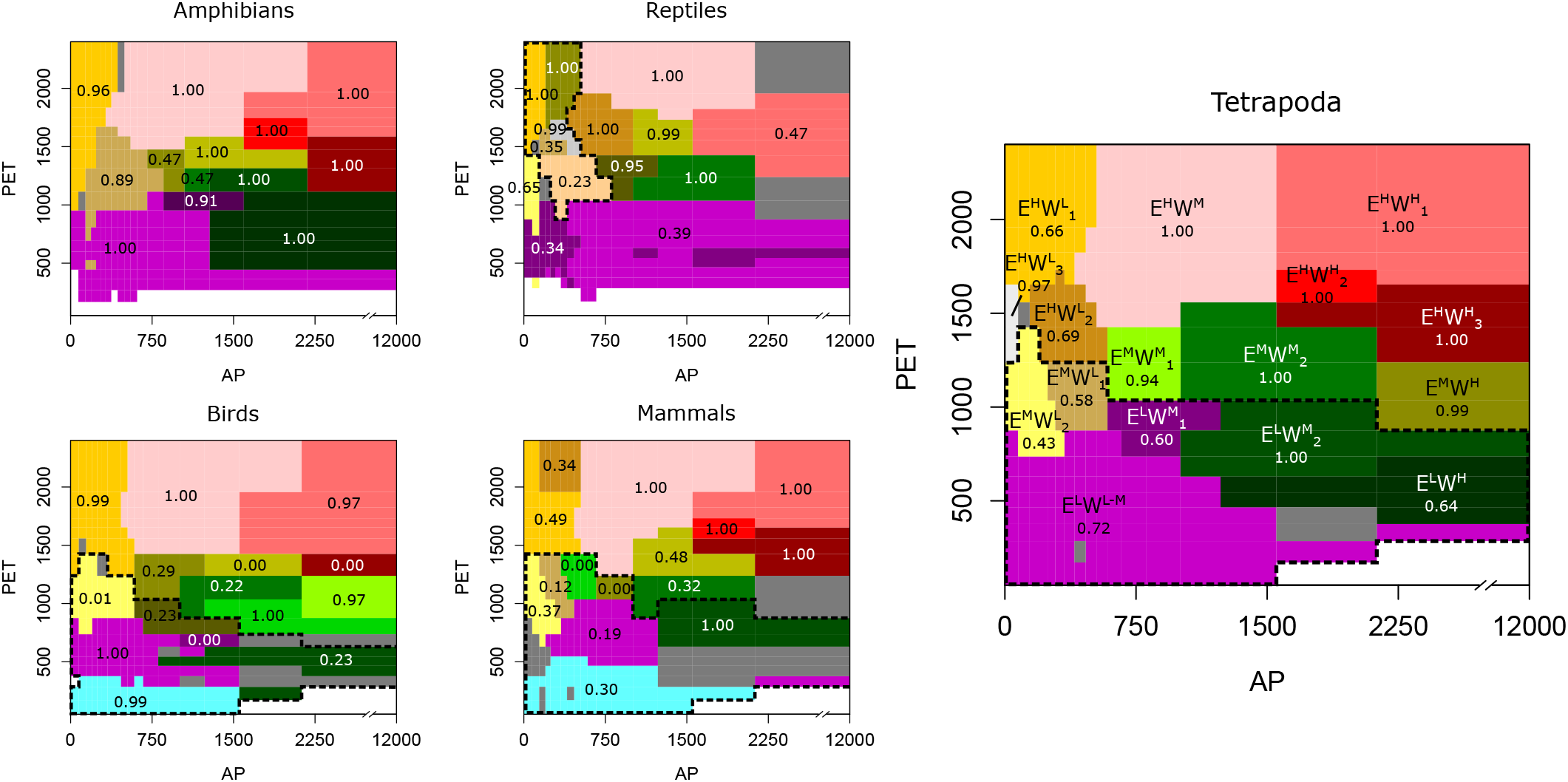
Tetrapoda niche domains across the climatic space. The climatic niche domains of each group shown across a space defined by potential evapotranspiration (PET) as a surrogate of energy and annual precipitation (AP) as a surrogate of water inputs. Tetrapoda superclass domains labelled so that E and W represent energy and water, respectively, and superscripts H, M and L mean high, medium and low, respectively. Numerical subscripts differentiate domains of similar climates. Bootstrap support between 0 and 1. The dotted line represents the domains at the highest hierarchical level. Domains formed of less than 50 species coloured in dark grey.

Since uncertainties related to the ranges of species exist, we employed a bootstrap and a significance clustering procedure [43, 44] to asses the domain robustness (Appendix S2). While several domains were well supported, we found that the niche domains corresponding to intermediate energy (between approximately 1000 and 1500 PET units; E^M^ climates in Fig. 2) and low to moderate water (up to approximately 800 m.m.l.l.; W^L^ to W^M^) were among the least supported. This robustness analysis shows that these niche domains are more challenging to classify.

### Tetrapoda vs Köppen’s climatic regions

With delineated niche domains, we studied the geographic location of their climatic conditions, the climatic regions in Fig. 1 and 3, which allowed for a more precise comparison between groups and Köppen’s regions. The similarities among the regions of Tetrapoda classes measured as Adjusted Mutual Information (AMI) ranged from 0.57 to 0.68, with mean AMI = 0.62 (Table S1). Moreover, the regions based on the niche domains of the superclass Tetrapoda were mostly congruent with the regions of its independent classes (mean AMI = 0.71, ranging from 0.66 to 0.77). Köppen’s regions were more dissimilar both to the regions of Tetrapoda (AMI = 0.44) and the ones of Tetrapoda classes (mean AMI = 0.44, ranging from 0.40 to 0.47).

**FIG. 3:**
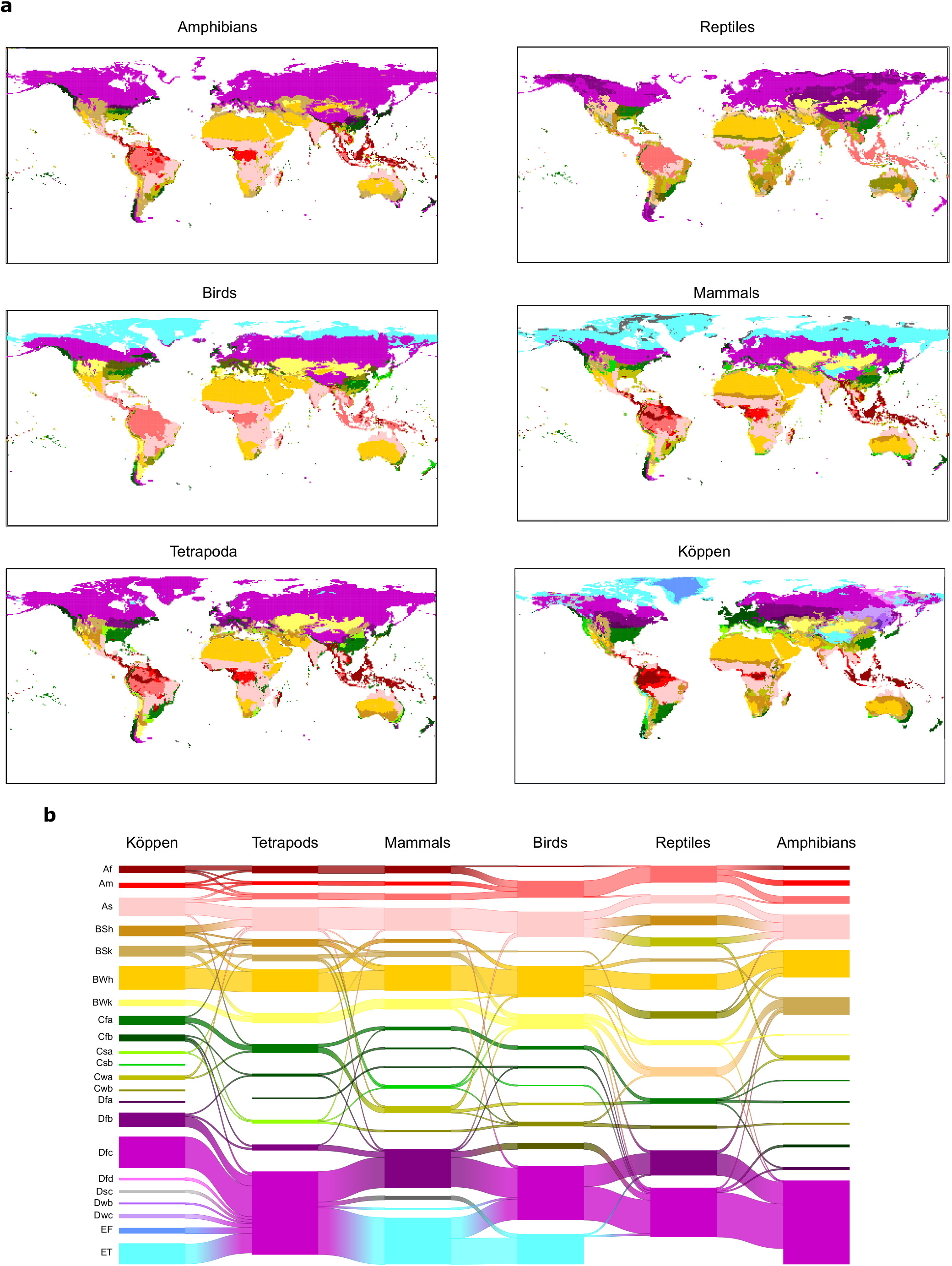
Tetrapoda groups and Köppen’s climatic regions are largely congruent. **a** Geographic location of Tetrapoda niche domains and Köppen’s climatic regions. **b** Alluvial diagram showing the similarities among the climatic regions. Colours according to Fig. 2.

Focusing on particular regions, we saw that climates
176 of high energy (E^H^) were consistent among groups and Köppen’s classification. Desert climates (high energy and low water, E^H^W^L^, BWh and BWk according to Köppen’s system) were the most similar across all classifications. Tropical savanna and steppe climates (high energy and medium water, E^H^W^M^, Aw and BSh respectively following Köppen) were also consistently defined, though both of these Köppen regions were often classified together in all groups but reptiles (Fig. 3). Similarly, Köppen’s tropical rainforest (Af) and tropical monsoon (Am) climates were for the most part well recovered. However, we found three different tropical-humid regions, each one mostly corresponding to one of the three largest masses of tropical rainforests: Amazonian, African and Southeast Asian rainforests; E^H^W^H^_1_, E^H^W^H^_2_ and E^H^W^H^_3_, respectively (Fig. 3). Regarding regions of low energy (continental, E, and polar climates, D, corresponding to the highest hierarchical level in Köppen’s system), we found a slightly higher level of disagreement between Köppen’s and Tetrapoda classifications (Fig. 3). Finally, temperate climates (medium energy E^M^) were the least congruent between groups and Köppen’s regions. Regions of medium energy were at the same time the least congruent among the different classifications and the least supported by the bootstrap analyses, suggesting that these climates impose less restrictive conditions.

### Climatic transition zones

A complete understanding of niche domains and their associated climatic regions entails exploring whether the domains have hard or diffuse transitions. Climatic conditions corresponding to diffuse transitions should present low specificity levels to the domain where they belong (Fig. 1). Our network approach allows to calculate this specificity by the dual classification of climatic bins and species into same niche domains (Fig. 1). We computed the specificity of each climatic bin as the ratio between the link weights of the species classified in the same domain and the total link weights [3, 45]. Then, we projected these values geographically. As expected, lower specificity values were in general associated with the boundaries of the climatic regions (Fig. 4a and S1). Our results also revealed that harsh conditions, such as desert and continental-polar climates (E^H^W^L^ and E^L^W^L^), present the highest specificity levels, regardless of the group (Fig. 4a and S1), reecting the dificulties to colonise these climates. Contrarily, temperate regions showed the lowest levels of specificity. These regions were also weakly supported in the bootstrap analysis; we found that bootstrap *p*-values and mean specificity were significantly correlated (stand. Glmm. coeff. 6.21; P < 0:001; *R*^2^ conditional = 0.29, see Material and Methods). Together with the higher variability of these regions across groups, this result further supports the idea that these climatic conditions could impose less restrictive conditions to Tetrapoda.

**FIG. 4:**
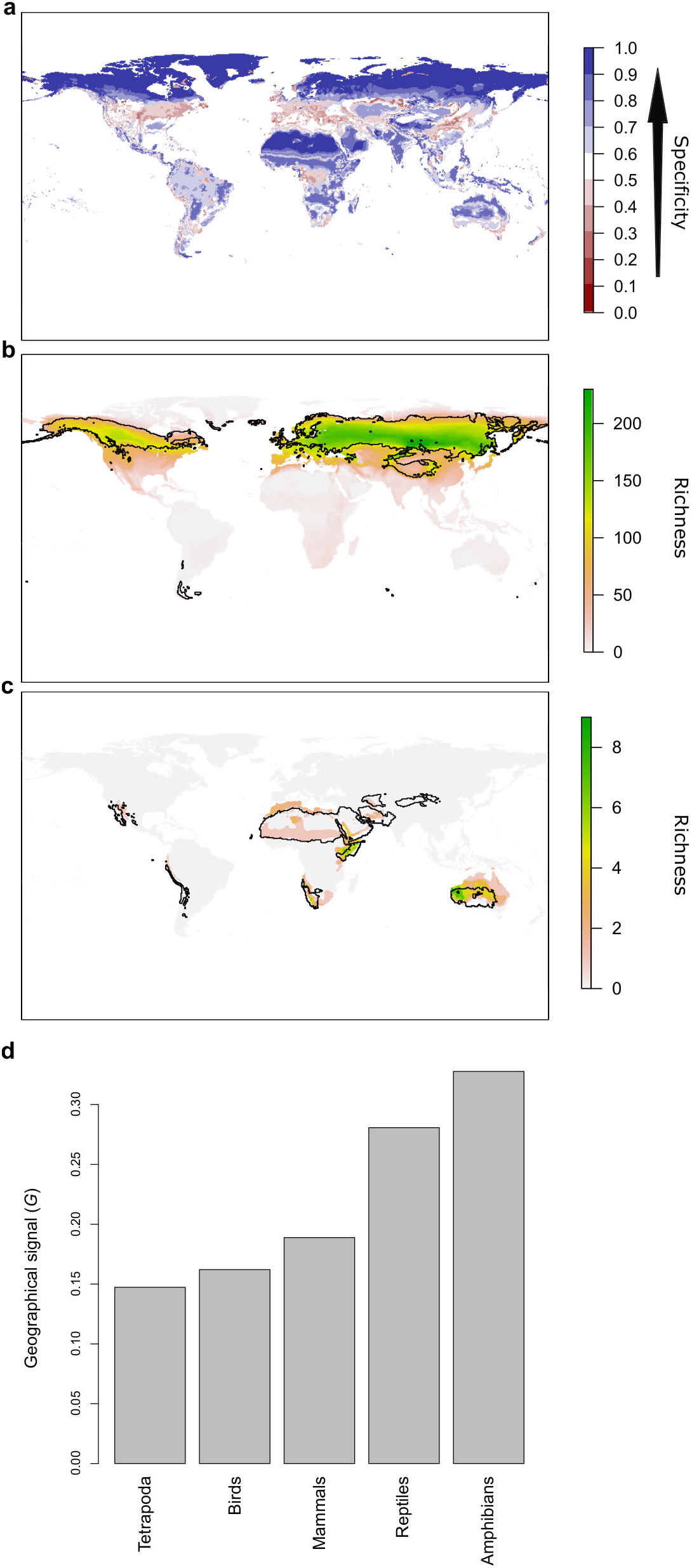
The geographic location of climatic domains and their associated species provide insights into the mechanism underlying the climatic regions. **a** Geographic projection of the specificity of climatic bins to their niche domain. **b** An example showing a bird’s niche domain with a low geographical signal. The distribution of the climatic conditions (black line) and the species (coloured richness values) belonging to the same niche domain were mostly congruent. **c** An example of an amphibian’s niche domain showing a high geographical signal, reflected in a substantial mismatch between the distribution of climatic conditions and species belonging to the same domain. **d** A quantitative approximation of the geographical signal, ranging between 0 and 1, for the different taxonomic groups (see Materials and Methods).

### Geographical signal in climatic regions

Historical and geographical processes can produce the detection of climates leading to specific species pools in some regions of the Earth but not in others. Thus, to explore for this geographical signal, we first compared the distribution of the climatic conditions and species grouped within the same niche domain. A geographic mismatch between species and climate distributions would point to portions of the climatic regions that are defined by species occurring in other geographic areas. Exploring these patterns for each niche domain revealed notable geographic agreement between species and climatic conditions of the same domain (Figs. 4b and S2-6). Nevertheless, we found some differences across groups and regions. More extreme climates showed larger mismatches between species and climates distributions. For instance, for all groups but reptiles, desert climate (E^H^W^L^) was mostly defined by species inhabiting Australia and to a lesser extent by species from the Namibian desert and The horn of Africa, with few or none species inhabiting the Sahara desert (Figs. 4c and S2-S6). Similarly, the northern climatic regions of amphibians and reptiles were defined by species at lower latitudes (Figs. S2-3). Approaching the geographical signal more quantitatively (see Material and Methods), we found a stronger signal for the worse dispersers amphibians and reptiles than for mammals and birds (Fig. 4d), suggesting that dispersal capabilities can contribute to the geographical signal in the niche domains. Finally, the Tetrapoda superclass showed the lowest geographical signal, which suggests that, beyond dispersion, an increased evolutionary time can reduce the geographical signal.

## DISCUSSION

We detected 16 climatic regions governing the distribution of Tetrapoda. Despite the substantial physiological and functional difierences among the groups, most of their niche domains and climatic regions are consistent. Some of these climatic regions resemble Köppen’s regions, which supports the idea that general climatic constraints organise the distribution of life on Earth.

While we found a high general congruence across groups, some niche domains and climatic regions were more consistent than others. In general, more extreme climates, such as arid or low-energy continental areas, were well defined in all groups. These climates also presented high levels of specificity, showing that species adapted to other climates have more dificulties to with stand these conditions. Both of these results suggest that extreme climates impose strong adaptive barriers [46, 47], even across distinctive evolutionary lineages.

Contrarily, milder climatic conditions, especially temperate climates, showed the lowest specificity, statistical support, and congruence across groups. These climates are more dificult to classify due to the overlap in the climatic space of species pools with different climatic optima. Two complementary reasons can explain this ambiguity: First, while we used two variables widely recognised to shape Tetrapoda distributions, alternative variables, such as seasonal changes of energy and precipitation [48], may also inuence species inhabiting temperate regions. Including these variables might help to further separate temperate species pools across the climatic space. Second, the climatic conditions of these domains may not prevent the colonisation of species with other realised optima or preferences, which would generate the observed overlap in the climatic space across milder conditions. Questions remain about the relative contribution of each factor.

We also found some domains that were well supported but unique for each group. These differences between groups seem to relate to the particular physiological adaptations of each group. For instance, homeothermic birds and mammals defined a region of low energy, consistent with Köppen’s polar climates, that reptiles and amphibians lacked. Similarly, reptiles, a group holding several species adapted to arid environments [36], defined some regions of low precipitation and high PET. Hence, despite the high similarities among groups, our results stress that caution is needed when generalising the climatic regions to other groups of organisms.

Beyond niche domains, our results also show differences in the geographical signal across groups. That amphibians – species with the lowest dispersal capacity – showed the highest geographical signal suggests that dispersal processes play an essential role: worse dis-perser species have more dificulties tracking their preferred climates [49], limiting the colonisation of disjoint areas with similar climates. Moreover, the Tetrapoda superclass shows the lowest geographical signal, which suggests that a increased evolutionary time can reduce this effect. Thus, evolutionary history – through the appearance of convergent adaptations to similar climates in different geographic regions[50] – may also inuence the geographical signal in niche domains. In any case, the ultimate causes and consequences of this signal require further attention. Why are some amphibians able to inhabit arid conditions in the Australian desert but not in the Sahara desert (Fig. S2)? Why can some reptiles withstand cold climates in and around the Himalayan mountains but not at the high latitudes of the northern hemisphere (Fig. S3)? These are some of the emerging undamental questions whose answers require historical biogeographical and evolutionary approaches.

Our results bring us closer to a definition of climatic regions that represent active factors for the organisation and evolution of life. Nevertheless, it would be interesting to improve some aspects in future studies. First, while we used a large number of species (about 26,000), they are taxonomically biased and only represent a small fraction of the terrestrial organisms. Similarly, we used two climatic variables widely known to affect the distribution and diversity patterns of animals and plants in general [4, 33], but other climatic variables might refine some of the least supported regions. Finally, our domains represent portions of the realised climatic niche space, and this space may be influenced by historical, geographical, and biotic factors beyond pure climate [3, 27, 51]. Although the geographical signal was rather low, identifying potential niches may also improve the accuracy of climatic regions. At the current pace of biological data accumulation and computational development, it is reasonable to expect that some of these limitations will soon be overcome. Meanwhile, the considerable congruence of several climatic regions across the studied groups and Köppen’s system provides confidence in their robustness. Hence, it is likely that using more and better data would not produce regions substantially different from those presented here.

Regardless of how generalisable the results are, the niche domains and their associated species pools and climatic regions can be used as a basis for ecological and evolutionary studies, as well as for conservation planning concerning Tetrapoda. Some of the many questions that the results reported here (data available in Appendix S3) can help to answer include : Are all the climatic regions similarly conserved and/or protected? Do the species forming each niche domain differ functionally or phylogenetically? Is the adaptation to niche domains evolutionary constrained? Do diversification, extinction or peciation rates differ among the species associated with different domains?

In conclusion, our data-driven climate classification reveals major climatic boundaries organising the distribution of life on Earth. Questions remain regarding the mechanism underlying differences between groups in the climatic regions and the geographical signal. Nevertheless, the regions that are consistent across groups can help answer questions in a diverse array of fields, including climatology, geography, ecology, evolution and conservation.

## MATERIAL AND METHODS

### Data

We obtained the distribution ranges of mammals and amphibians from The IUCN Red List of Threatened Species [21], of birds from Bird species distribution maps of the world [22] and of reptiles from ref. [23]. We included only the native range of terrestrial species in the analyses in all instances. In the case of birds, we only used the breeding ranges. Moreover, since there is a higher uncertainty when determining the realised niches of narrow-ranging species [52], we removed the species whose ranges were less than 5 grid cells of 0.5 degrees. After this cleaning of the data, we used 3657 amphibians, 7204 reptiles, 4574 mammals and 10684 birds, for a total of 26119 Tetrapoda species.

We approximated the species’ Grinnellian niches[51] with two climatic variables that represent energy and water inputs. While we could have used several other variables, we chose energy and water since they best explain climatic effects on species distributions [4]. As surrogates of energy and water inputs, we used mean annual potential evapotranspiration (PET) and annual precipitation, respectively. Both variables have been shown to be important factors for Tetrapoda species distributions [33–35]. Moreover, they have also been used in previous climate classifications [18] and are regularly used to derive other drivers of species distributions such as the UNEP aridity index [53, 54]. We obtained PET from ref. [25] and annual precipitation from ref. [24], both at a 0.08° resolution. Finally, we obtained Köppen’s climatic regions from refs. [9, 55].

### Niche characterisation

We characterised the realised climatic niche of each species using an approach similar to the one proposed in ref.[26]. We divided the climatic space formed by PET and annual precipitation into bins and calculated the proportion of occurrences a given species has in each climatic bin. Both the shape of the divisions and the number of divisions of each climatic axis affect the result. For instance, dividing the axis into regular intervals can destroy critical information if the climatic values more restrictive are skewed toward any extreme of the distribution or if the climatic values are represented non-uniformly across the globe (as for annual precipitation, Fig. S7). Also dividing the space into too few intervals destroys information, whereas using too many divisions can generate niche domains with only a few species. To overcome the first issue, we divided the axes in quantiles based on the distribution of climatic values across the Earth. By doing so, we obtained an almost uniformly divided PET axis (Fig. S7). Contrarily, the number of divisions of the annual precipitation axis was skewed towards low values, which resulted in a higher resolution over the presumably more relevant low-precipitation conditions (Fig. S7). To solve the second issue, we selected the lowest number of divisions that maximised the gain in information (see Appendix S1). The optimal number of axis divisions was 17 in all cases but amphibians, where it was 18 (Fig. S8).

Next we accounted for potential commission errors, which may affect the different climates a species experiences. Specifically, range maps can overestimate the area occupied by a species, which directly inuences the niche characterisation [56]. Extracting the climatic values that a species range covers from a high-resolution climatic raster (such as 0.08°) may reduce commission errors at the borders of the species range, but increases this error otherwise. Extracting climatic values from a coarser raster can reduce the inuence of commission errors over the areas inside of a range but increases them over the borders. To alleviate the effects of these potential errors, we first extracted the climatic values from the high-resolution rasters (0.08°). Then, we computed the average climatic values among selected raster pixels located within cells of 0.5 degrees. In this way, we reduced the effects of commission errors both at the borders of and inside species ranges. Moreover, we also conducted a bootstrap significance test that takes uncertainty of species ranges into account (see below).

### Niche domains and climatic regions identification

We employed a network community detection approach to identify the niche domains and the species mainly associated with them. For each group of species, we first generated a weighted bipartite network where species and climatic bins formed the disjoint sets of nodes, and the proportion of occurrences of species in intervals of the climatic values corresponding to the climatic bins formed the weighted links. We then used the hierarchical version of the community detection algorithm known as Infomap [41, 42] to identify the niche domains. We ran the algorithm 1000 times, selecting the network partition with the best quality.

To consider the uncertainty associated with both the species ranges and the community detection, we conducted a bootstrap analysis. For each species, we resampled with replacement from the distribution of climatic values within species ranges at a resolution of 0.08°. We veraged climatic values laying within 0.5° cells and calculated the proportion of occurrences in each climatic bin. With resampled data from all species, we generated a bootstrapped network and ran Infomap 1000 times using this network. Given the high computational cost of this analysis, we only generated 100 bootstrap networks. We followed the approach proposed in ref. [44] to calculate the support of the niche domains. For each identified domain, we calculated the proportion of bootstrap networks with a domain more similar than Jaccard index 0.5 [44].

With obtained niche domains, we detected the climatic regions by identifying areas across the Earth’s surface that hold the climatic conditions grouped within each niche domain. Finally, to compare climatic regions across Tetrapoda groups and with Koöppe’s classification, we calculated the adjusted mutual information (AMI) [57].

### Climatic transition zones

The joint classification of climatic bins into domains and the species most associated to them allowed us to calculate the specificity of the bins to the domain where they belong, which indicates zones of transitions between domains (Fig. 1). That is, a bin acting as a transition between two domains should contain species from both domain and, therefore, a low specificity to the domain where it is classified [3, 45]. To consider the link weights, we calculated this specificity 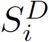 of a climatic bin *i* in domain *D* as the sum of link weights *w*_*i*;*j*_ of the species *j* present in the bin and also belonging to the same domain as the bin, divided by the sum of link weights of all the species present in the bin, such that

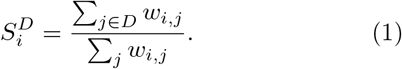

This index is 1 when the bin has only species of the same domain as the bin, and tends to 0 otherwise. We then projected these specificity values into the geographic pace by assigning these values to the geographical raster cells *q* that hold the climatic conditions represented by the bins, thus obtaining the projected specificity 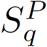. Finally, we explored the relationship between average SD and bootstrap support. We fitted a logistic GLMM of bootstrap *p*-values as function of mean *S*^*D*^ as fixed term and the taxonomic group as a random intercept term. GLMM was conducted using the lme4 [58] package in R [59].

### Geographical signal

To investigate the geographical signal, we first observed the match between the geographic location of the species and the climatic conditions associated with the corresponding niche domain. Then, we quantified the geographical signal by comparing the geographically projected specificity *S*^*P*^ with a measure of specificity based on the actual pool of species co-occurring geographically. That is, the specificity of a climatic bin *S*^*D*^ is based on the species that co-occur in the climatic space and then it is projected geographically to obtain *S*^*P*^ (see above). Hence, *S*^*P*^ does not considered the actual pool of species co-occurring in the geographic space. In case of a large geographical signal, we would expect large differences between the species co-occurring in the climatic and geographic spaces. For instance, the geographic mismatch between species and climates belonging to the same domain is produced by species co-occurring in a given portion of the climatic space but not in all geographical areas with the climate represented in such portion of the climatic space. In this sense, in case of geographical signal we would expect differences between the projected specificity *S*^*P*^ and a value of specificity based on the species pool occurring in given geographical areas, for short the ctual specificity *S*^*A*^. A higher actual specificity than the projected indicates areas that host most of the species associated with a niche domain, while the opposite indicates areas not, or only scarcely, colonised by these species.

Using Eq. 1, we calculated the actual specificity of a geographical raster cell *q*, whose corresponding climatic bin i is in domain *D*, as the ratio between the link weights of species in raster cell *q* that belong to its associated domain and the total link weights of species in *q*,

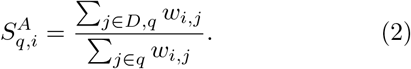

To calculate the geographical signal *G*, we computed the projected and actual specificity for each climatic raster cell *q* at 0.5° resolution. Then, we calculated the verage differences between projected and actual specificity in absolute terms, such that

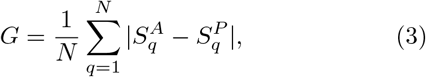

where *N* is the total number of raster cells. This index is 0 when there is no geographical signal and tends to 1 for high signals.

## Competing interests

The authors declare no competing interests.

## Author contribution

J.C. and M.N. conceived the ideas with inputs from all authors; J.C. analysed the data with assistance from all authors; J.C. wrote the manuscript in collaboration with all authors.

## Acknowledgements

We are thankful to Andrea Briega and Miguel Á. Rodríguez for discussion on early ideas. We are very grateful to Fernanda Alves-Martins, Rafaél Molina-Venegas, Cristina Roquillo and Rubén Bernardo-Madrid for critical reviews. J.C. is supported by the Carl Tryggers Foundation for Scientific Research (CTS 16:384).

