## Appendices S1-S2 for "Regularities in species niches reveal the World’s climatic regions"

Joaquín Calatayud, Magnus Neuman, Alexis Rojas, Anton Eriksson, and Martin Rosvall
*Integrated Science Lab, Department of Physics, Umeå University.*
(Dated: November 21, 2019)

### APPENDIX S1. SELECTING THE NUMBER OF BINS

Dividing the climatic space into an optimal number of bins is critical because it can affect subsequent classifications. On the one hand, an insufficient number of bins may destroy information. For instance, in the extreme case of using only one bin, all the species will belong to exactly the same climatic niche. On the other hand, an excess of bins may lead to niche domains with only a few specialist species, which are not suitable for a global climate classification. The optimal value is in between these extremes. Intuitively, the differences in the species' climatic niches increase with the number of bins. Yet, when the bins are sufficiently many to capture the main differences among species, increasing the number of bins will not increase such differences. Thus, when the number of bins are enough to describe the main information of the system, more bins will not significantly increase the information contained in the species niches. Therefore, when the number of bins already captures the main information, the slope of the relationship between the differences in species' niches and the number of bins becomes less pronounced.

To measure the average differences between the niches, we used the well-known Jensen-Shannon divergence (JSD). Assuming that each species niche is a probability distribution $P_i$ across the discretised climatic niche space,

$$JSD = H\left(\sum_{i=1}^{n} \frac{P_i}{n}\right) - \sum_{i=1}^{n} \frac{H(P_i)}{n}, \qquad (1)$$

where $H(P)$ is the Shannon Entropy for the species niche $P$ and $n$ is the number of species. The JSD is bounded between 0, when the probability distributions are equal, and $\log_2(n)$, when the distributions are completely dissimilar. To make results comparable we normalised JSD by $\log_2(n)$.

We discretised the climatic space by dividing PET and annual precipitation in quantiles, ranging from 5 to 50 and increasing in steps of 5. For each division, we computed the proportion of observations of each species in each bin and calculated the JSD using all species. Finally, we fitted a piece-wise regression of the increments of JSD ($\Delta JSD$) as a function of the number of divisions (Fig. S8).

### APPENDIX S2. SUPPLEMENTARY TABLES AND FIGURES

TABLE S1: Similarity of climatic regions measured using AMI.

| | **Koppen** | **Tetrapoda** | **Amphibians** | **Reptiles** | **Birds** |
|---|---|---|---|---|---|
| **Tetrapoda** | 0.442 | - | - | - | - |
| **Amphibians** | 0.403 | 0.660 | - | - | - |
| **Reptiles** | 0.449 | 0.702 | 0.581 | - | - |
| **Birds** | 0.453 | 0.771 | 0.607 | 0.637 | - |
| **Mammals** | 0.475 | 0.723 | 0.574 | 0.663 | 0.685 |

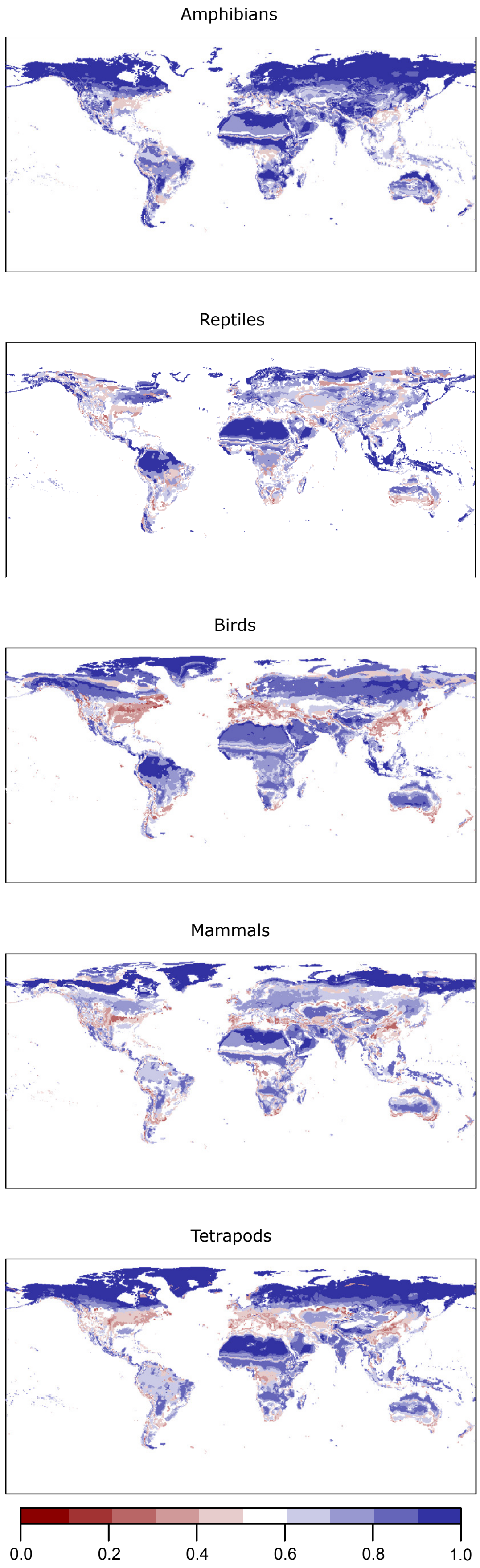


FIG. S1: Geographic projection of the specificity of the climatic bins ($S^P$) for the studied groups.

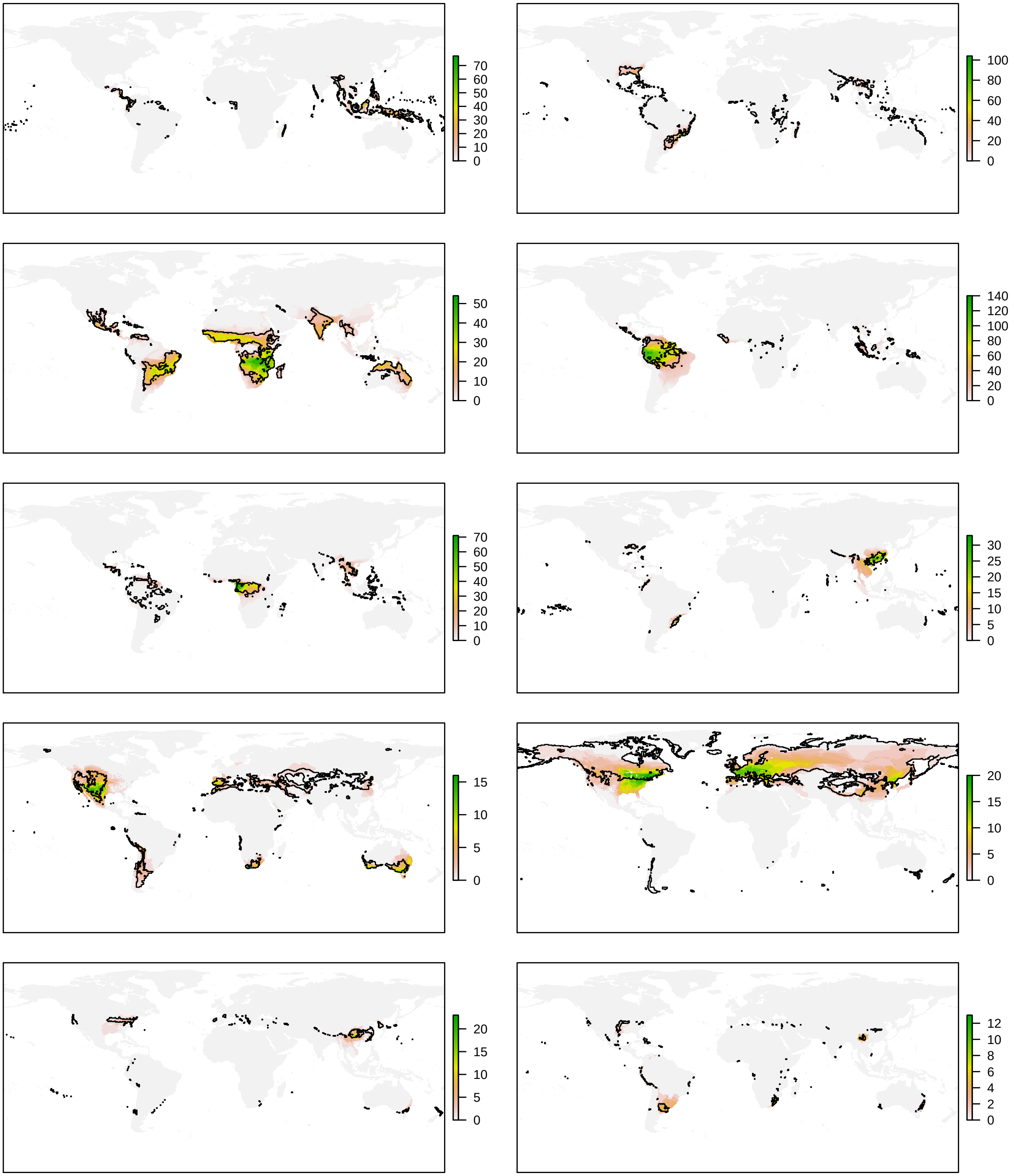


FIG. S2: Continues on the next page.

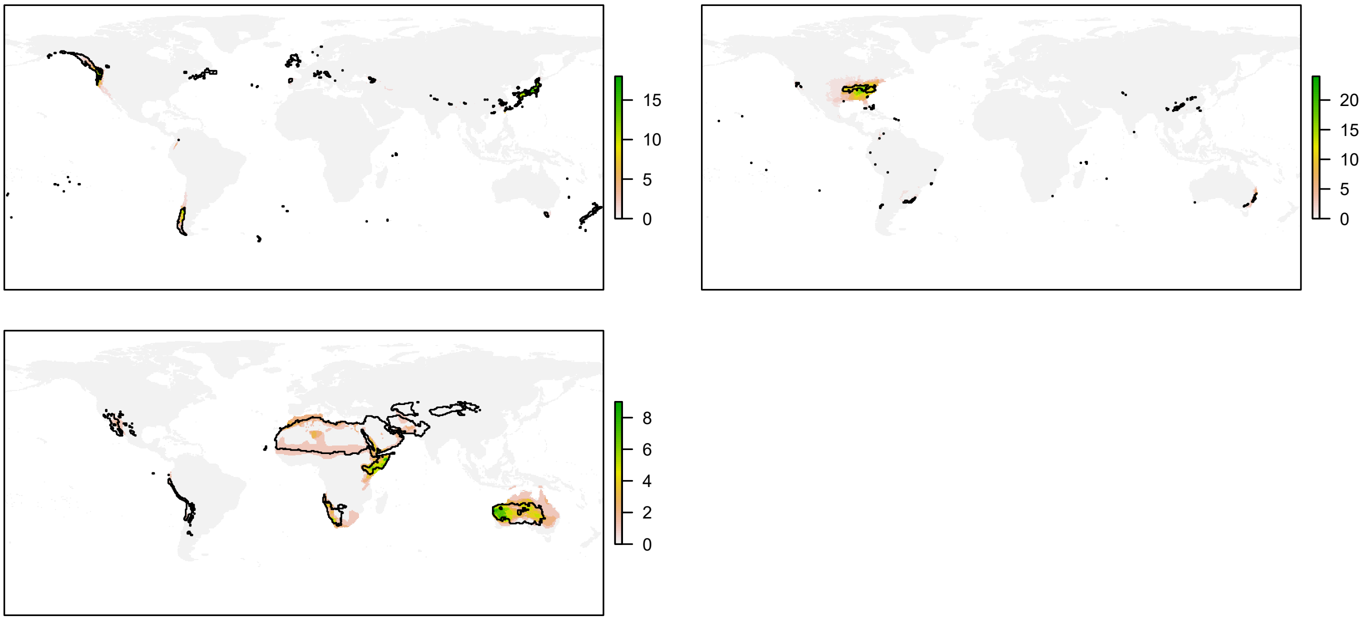


FIG. S2: The distribution of the climatic conditions (black line) and the species (coloured richness values) belonging to the same niche domains of amphibians.

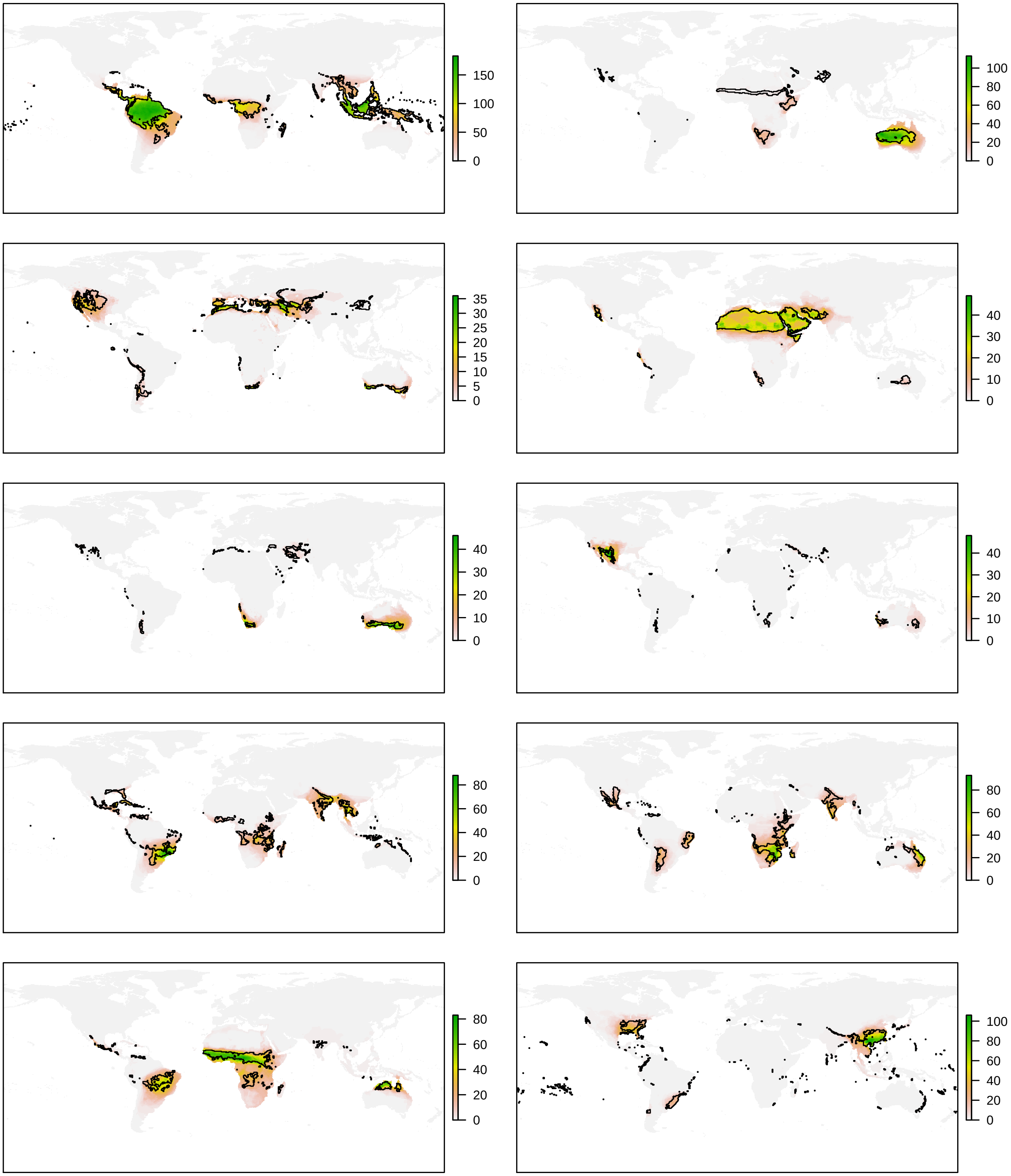


FIG. S3: Continues in next page.

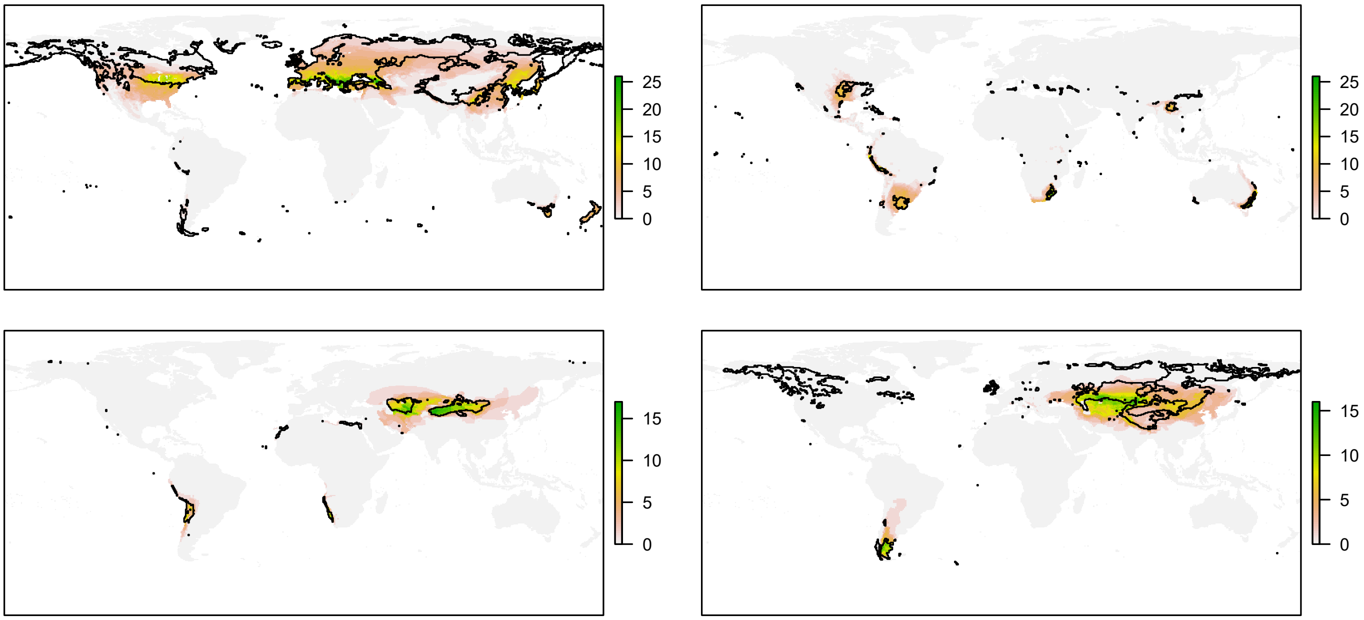


FIG. S3: The distribution of the climatic conditions (black line) and the species (coloured richness values) belonging to the same niche domains of reptiles.

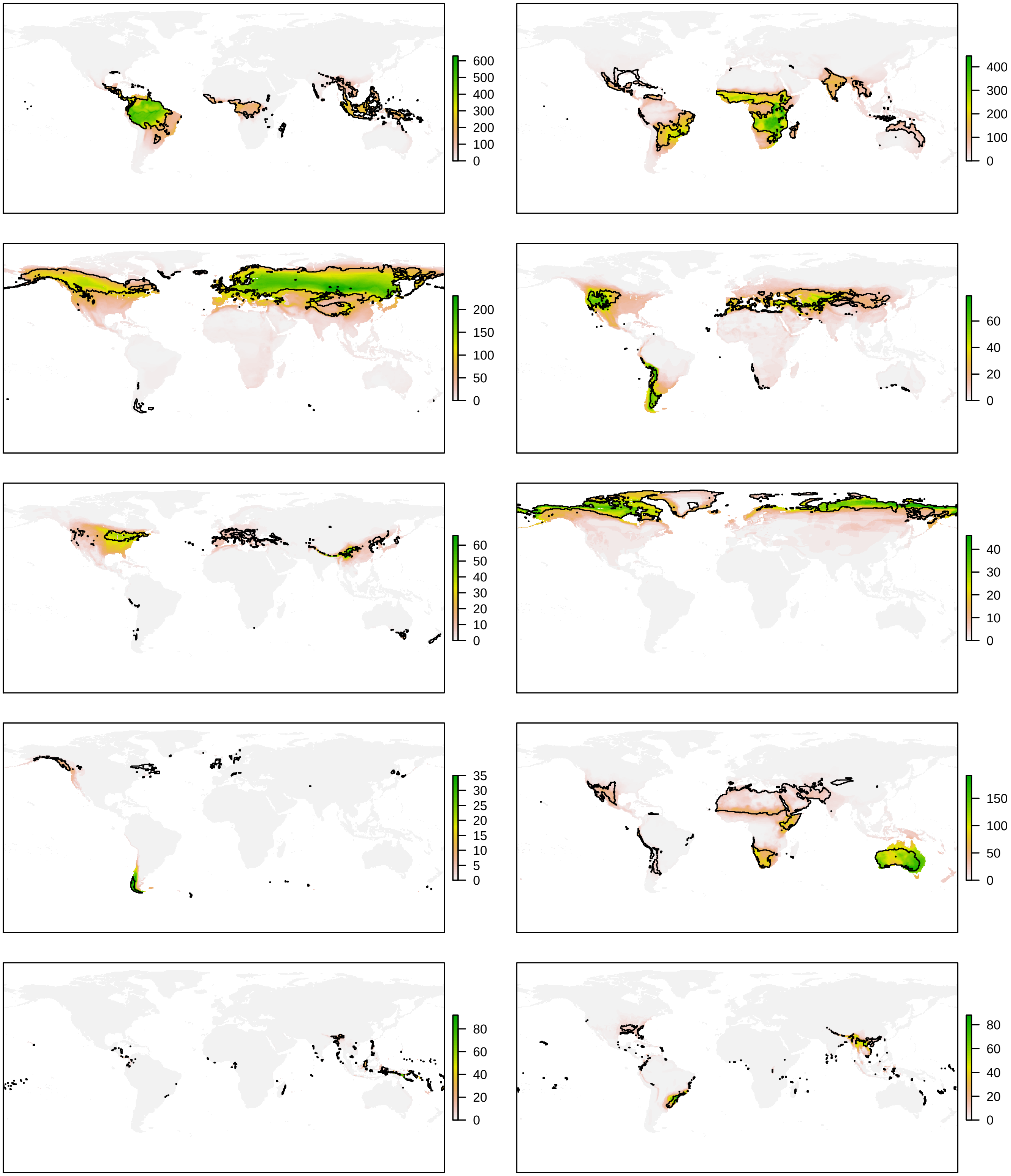


FIG. S4: Continues in next page.

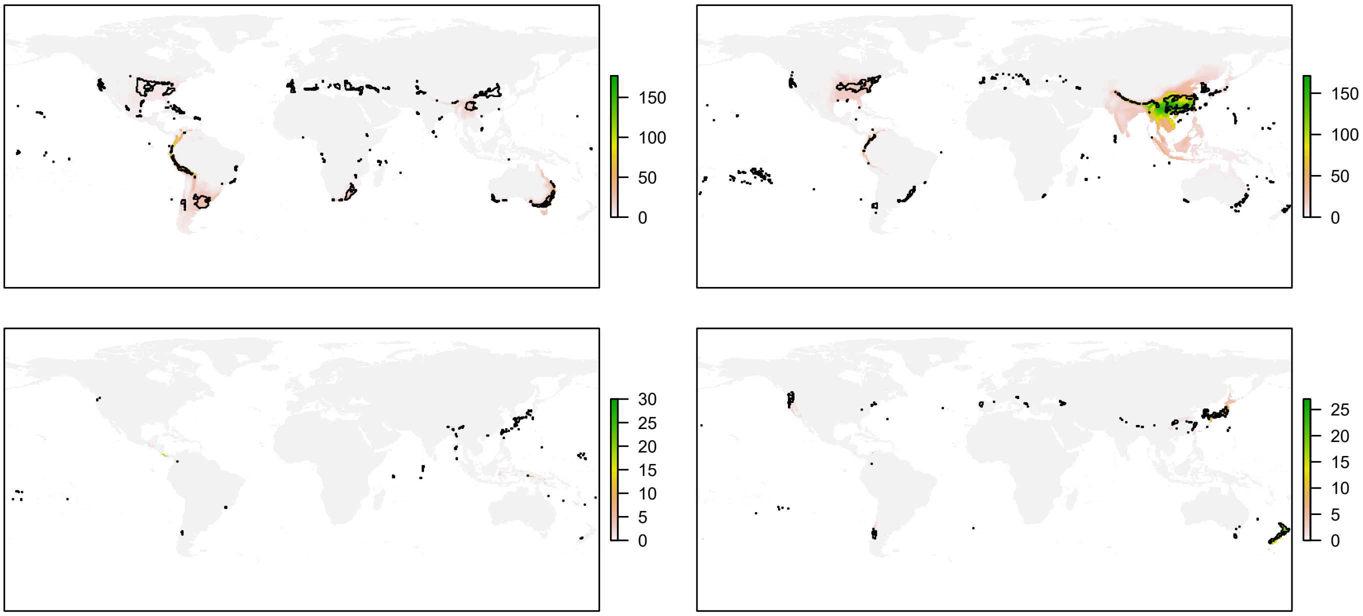


FIG. S4: The distribution of the climatic conditions (black line) and the species (coloured richness values) belonging to the same niche domains of birds.

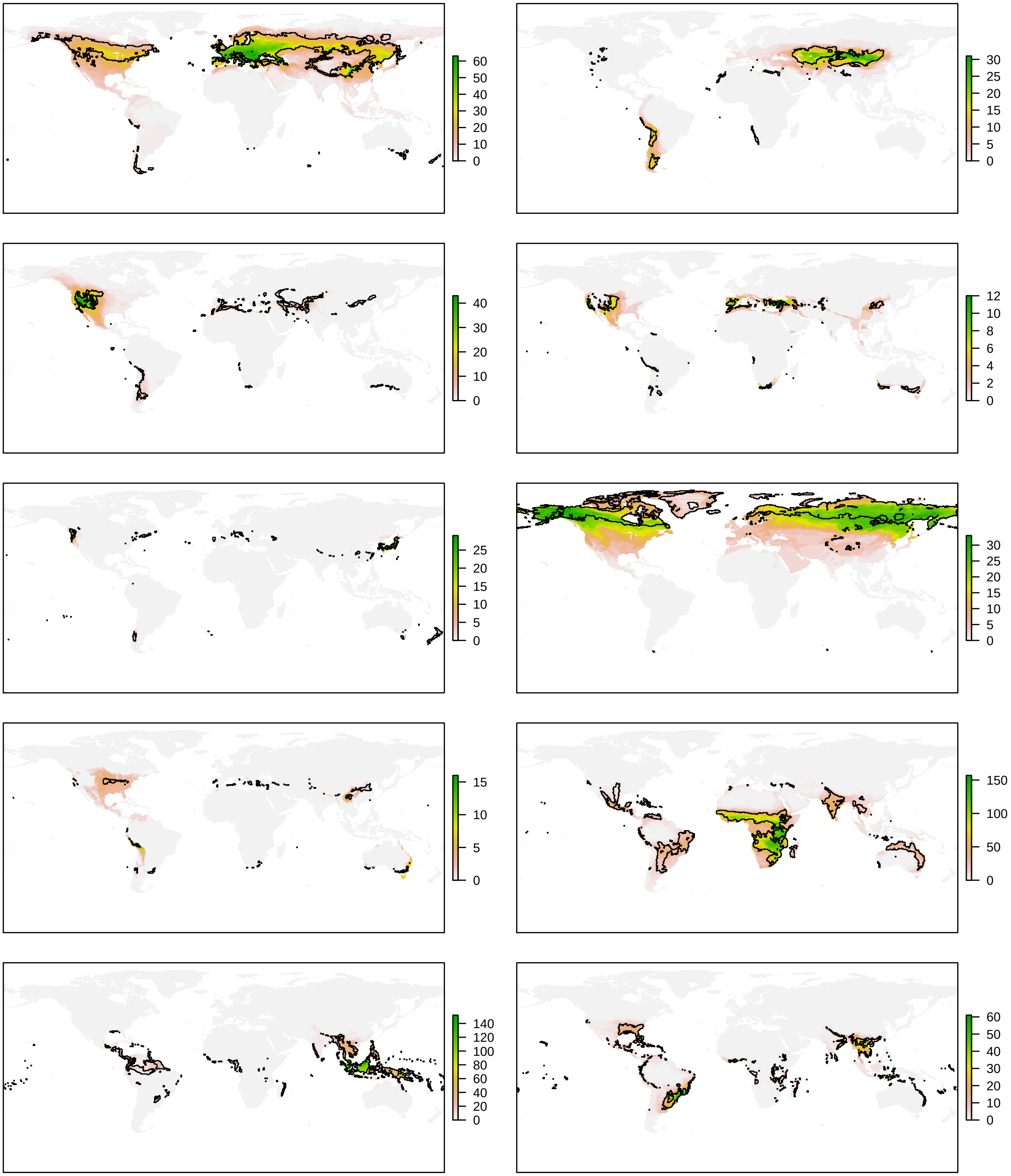


FIG. S5: Continues in next page.

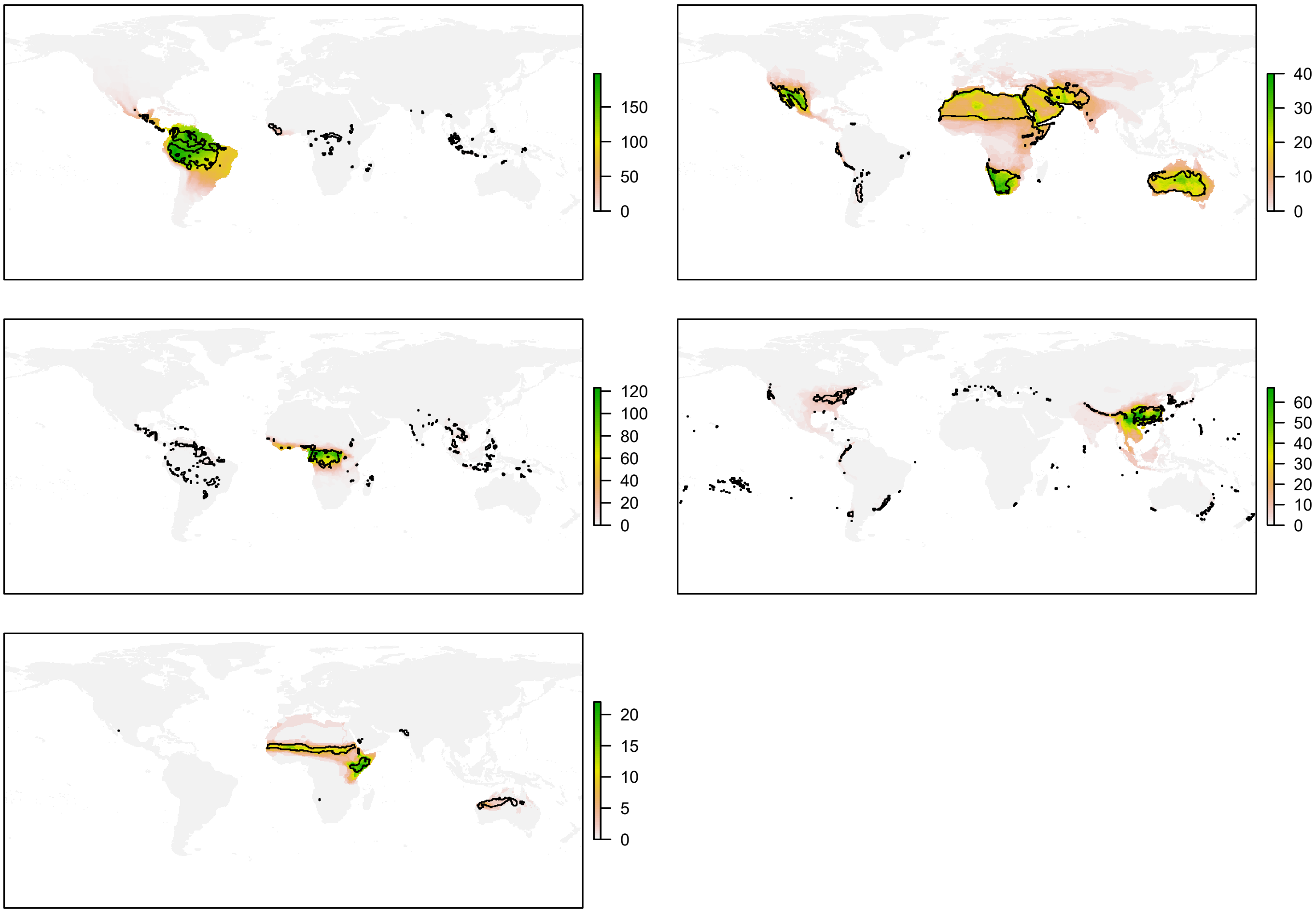


FIG. S5: The distribution of the climatic conditions (black line) and the species (coloured richness values) belonging to the same niche domains of mammals.

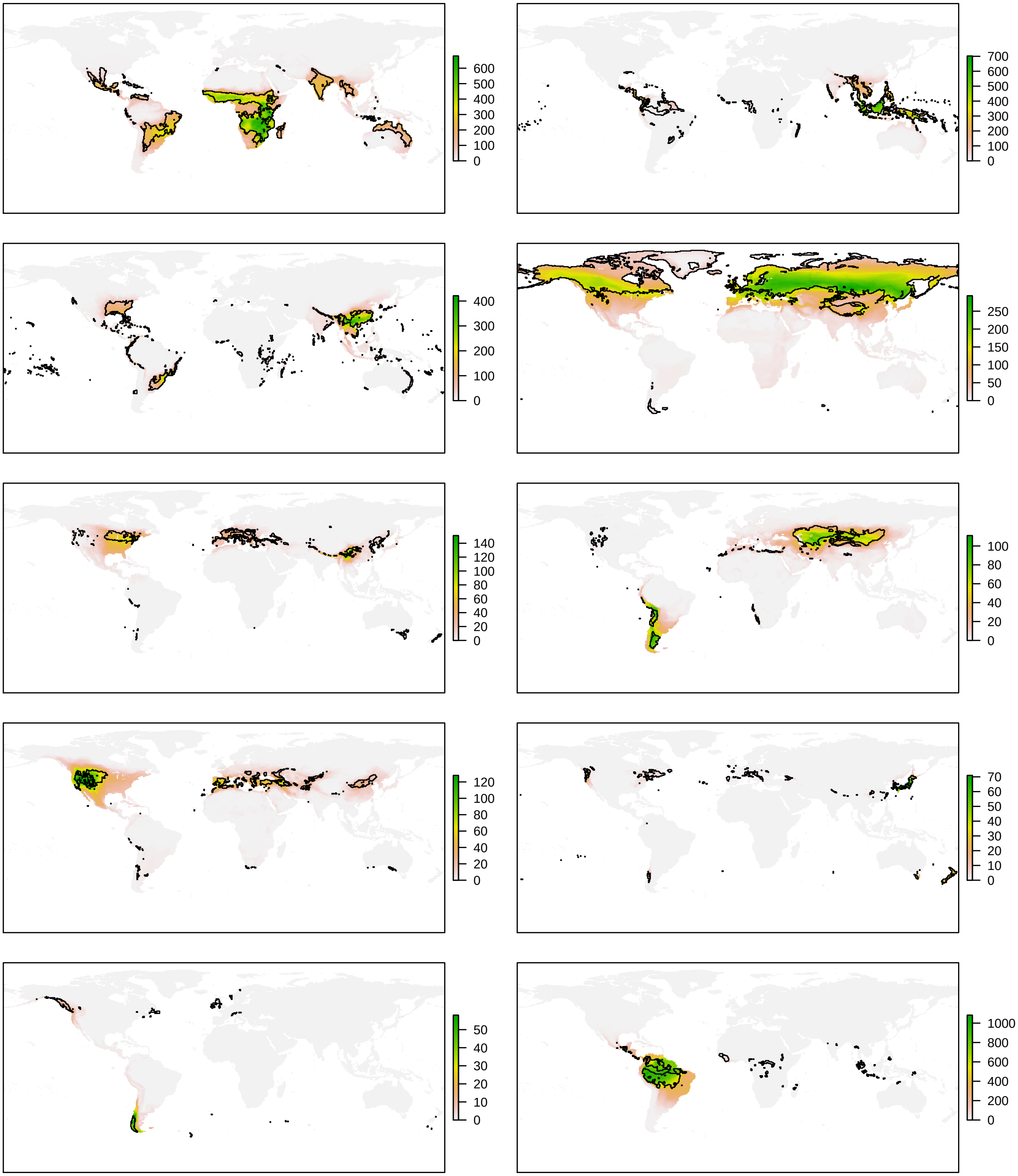


FIG. S6: Continues in next page.

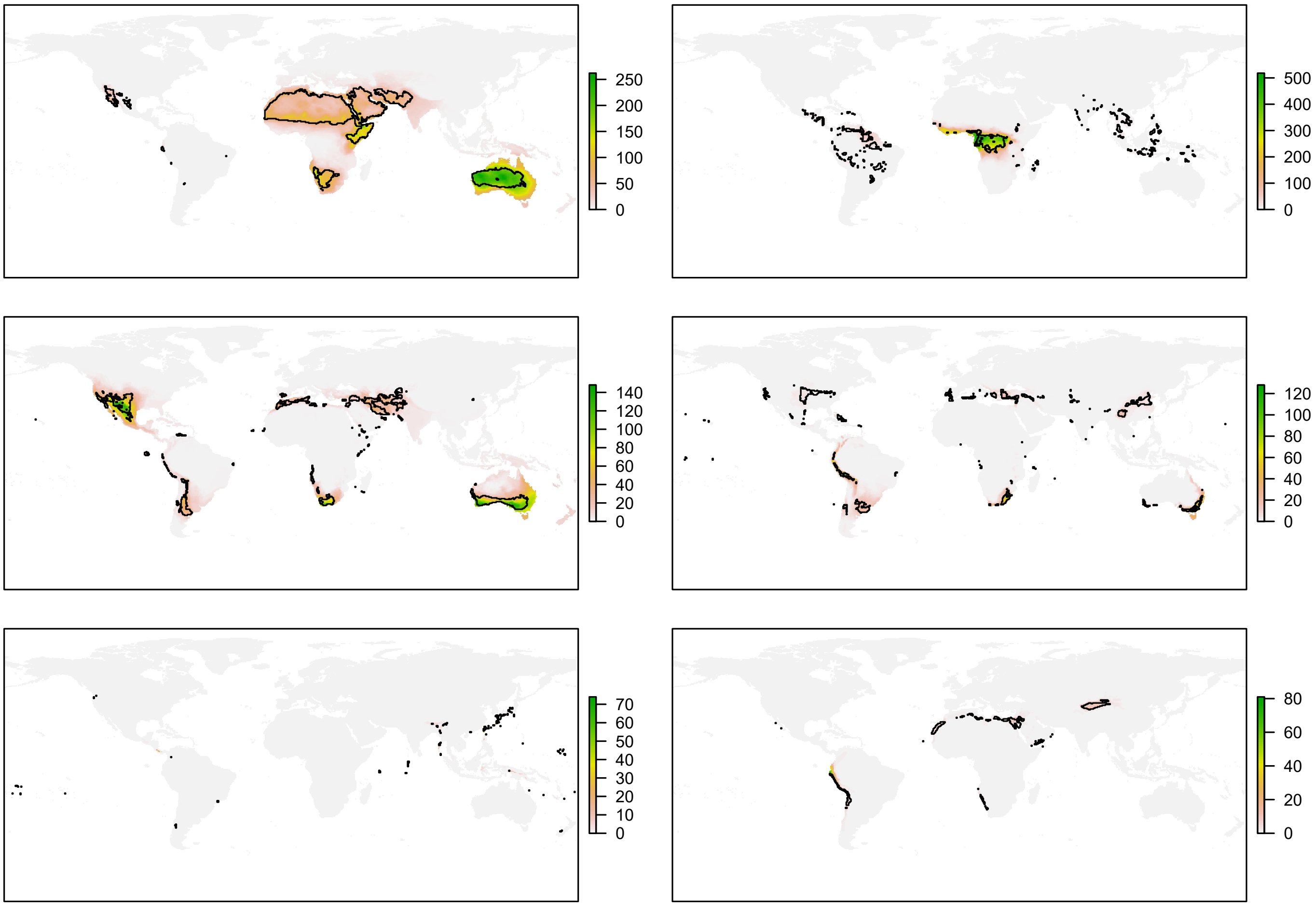


FIG. S6: The distribution of the climatic conditions (black line) and the species (coloured richness values) belonging to the same niche domains of tetrapods.

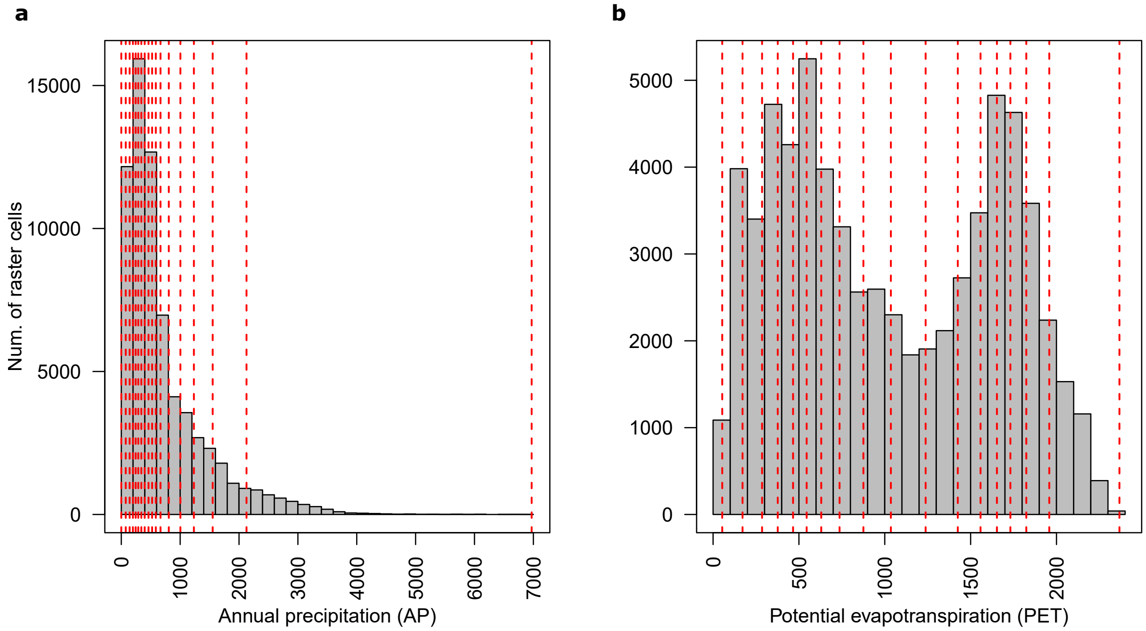


FIG. S7: Distribution of **a.** annual precipitation and **b.** potential evapotranspiration values. 17 divisions of each climatic variable are presented with red dotted lines.

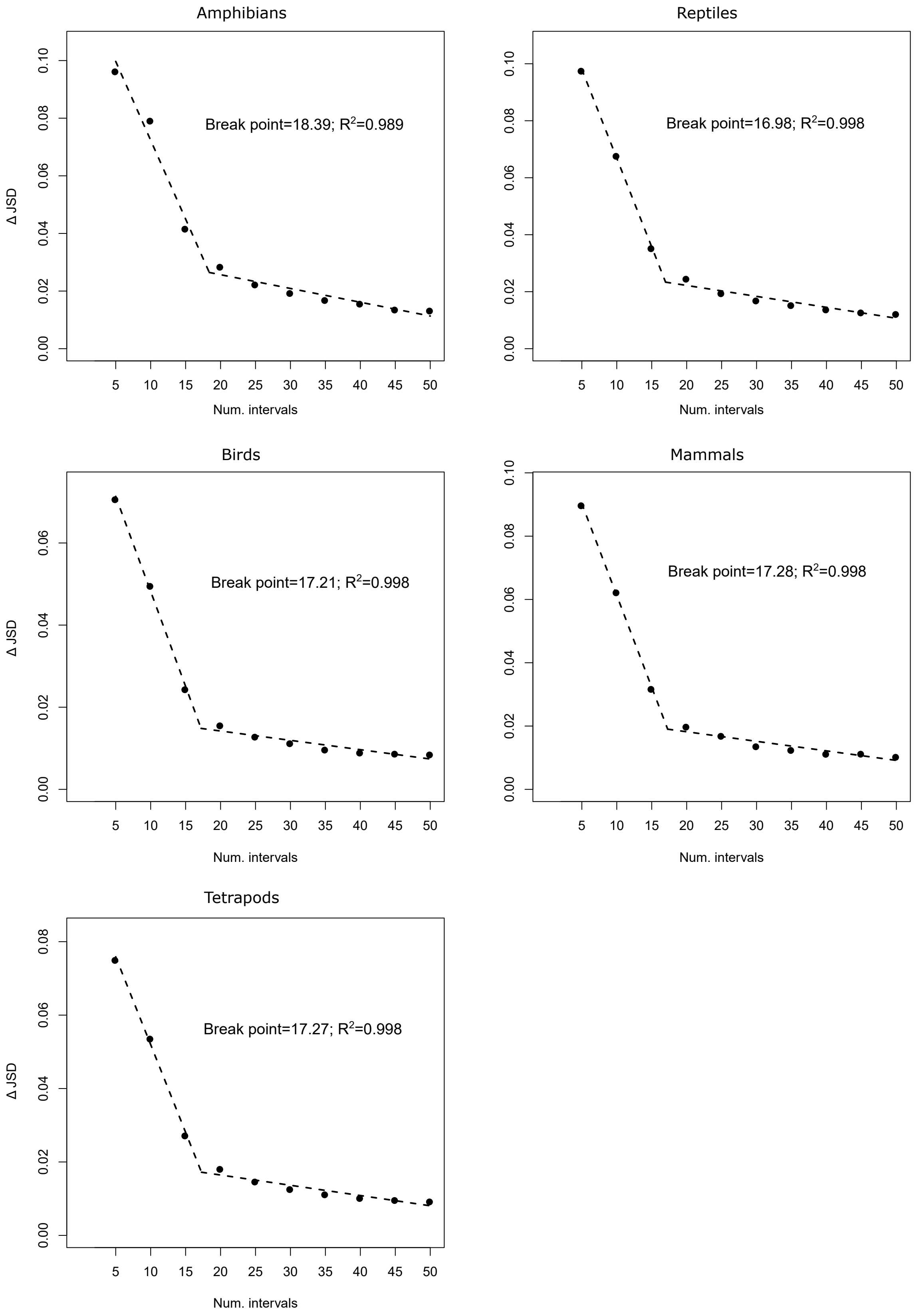


FIG. S8: Predictions of piecewise regression of the increment in JSD ($\Delta JSD$) as a function of the number of divisions in the variables defining the climatic space of all studied groups. The breakpoints and the coefficient of determinations are also provided.
